# Optimal balancing of xylem efficiency and safety explains plant vulnerability to drought

**DOI:** 10.1101/2022.05.16.491812

**Authors:** Oskar Franklin, Peter Fransson, Florian Hofhansl, Jaideep Joshi

## Abstract

- In vast areas of the world, the growth of forests and vegetation is water-limited and plant survival depends on the ability to avoid catastrophic hydraulic failure. Therefore, it is remarkable that plants take high hydraulic risks by operating at water potentials (*ψ*) that induce partial failure of the water conduits (xylem). Here we present an eco-evolutionary optimality principle for xylem conduit design that explains this phenomenon.
- Based on the hypothesis that conductive efficiency and safety are optimally co-adapted to the environment, we derive a simple relationship between the intrinsic tolerance to negative water potential (*ψ*_50_) and the environmentally dependent minimum xylem *ψ*.
- This relationship is constrained by a physiological tradeoff between xylem conductivity and safety, which is relatively strong at the level of individual conduits although it may be weak at the whole sapwood level. The model explains observed variation in *ψ*_50_ both across a large number of species, and along the xylem path in two species. The larger hydraulic safety margin in gymnosperms compared to angiosperms is explained as an adaptation to the gymnosperms’ lower capacity to recover from conductivity loss.
- The constant xylem safety factor provides a powerful principle for simplifying and improving plant and vegetation models.

## Introduction

The hydraulic properties of plants constrain their ability to grow and survive in different environments. Therefore, a solid understanding of these constraints is essential for accurate prediction of vegetation responses to droughts and other environmental changes. In stems, the primary function of the xylem is to transport water through the stem, driven by the difference in water potential between the base and the top. The conductivity of the xylem increases steeply with the conduit diameter as long as it is not hampered by cavitation, or embolism, which happens when water potential is too low (too highly negative). The sensitivity to cavitation is often measured in terms of the water potential at which conductivity is reduced by 50%, *ψ_50_*. Across a large number of species, *ψ_50_* is, on average, close to the minimum midday water potential (*ψ_min_*) experienced by each species, implying a remarkably small hydraulic safety margin (*ψ_50_ - ψ_min_*) (Choat *et al*., 2012). The safety margin is lower for angiosperms than gymnosperms and is smallest in wet sites with high *ψ_min_*. However, the reason for this apparently risky strategy and its variation with climate are not yet fully understood, despite the importance for our understanding of plant responses to expected climate changes and droughts (Venturas *et al*., 2017). Here we aim to explain the variation in *ψ_50_* through an eco-evolutionary-optimality perspective.

Eco-evolutionary optimality (EEO) approaches have been used to model a wide range of plant traits and processes (Franklin *et al*., 2020), including plant stomatal hydraulic regulation (Hölttä *et al*., 2011; Wolf *et al*., 2016; Anderegg *et al*., 2018; Wang *et al*., 2020), vascular network structure (McCulloh *et al*., 2003; Savage *et al*., 2010; Koçillari *et al*., 2021), and leaf hydraulic and photosynthetic traits (Deans *et al*.,2020). Xylem efficiency and safety in relation to growth environment has only rarely been addressed from an EEO perspective, perhaps due to a lack of consensus on the xylem costs and benefits. For example (Manzoni *et al*., 2013) used an EEO approach based on the principle that transpiration is maximized, which contrasts to the assumption in most other models that transpiration is a cost (Wang *et al*., 2017). Furthermore, although a tradeoff balancing xylem conductivity (benefit) and vulnerability (cost) is expected to constrain optimal xylem function, recent studies suggest that this tradeoff is weak (Gleason *et al*., 2016; Sanchez-Martinez *et al*., 2020; Liu *et al*., 2021). Nevertheless, such a tradeoff is invoked in the EEO-based widened pipe model explaining the stem tip to base widening of conduits (Koçillari *et al*., 2021).

Here we first show that the xylem efficiency-safety tradeoff is important at the level of conduits, even though it is weaker at the level of whole sapwood tissue. Then, we combine this tradeoff with an EEO hypothesis which states that plants maximize the combination of xylem conductive capacity per unit conduit biomass and tolerance to low *ψ*. Since conductivity loss is not always reversible (Anderegg *et al*., 2013; Pellizzari *et al*., 2016) we further extend the vulnerability concept to account for accumulation of irreversible conductivity loss. Our first-principles EEO-based model successfully explains the globally observed relationship between *ψ_50_* and the environmentally dependent *ψ_min_*, and how it differs between angiosperms and gymnosperms.

## Model description and results

### The conduit conductivity – safety tradeoff

The xylem can be seen as a network of interconnected conduits (vessels or tracheids). Xylem conductivity and safety (tolerance to negative water potential, *ψ_50_*) depends on both the conduits themselves and their interconnections, the pits and end-walls, and how the conduits are spatially clustered (Lens *et al*., 2011). The hydraulically weighted conduit diameter (*D*) is an important determinant of conductivity due to the strong effect on fluid dynamics (Hacke *et al*., 2017), which is also reflected in the ubiquitous tapering of conduits with stem height which serves to minimize the increase in resistance with path length (West *et al*., 1999). Following the Hagen–Poiseuille equation, the hydraulic conductivity (maximal conductive capacity, *K*) of conduits increases with the 4^th^ power of the conduit diameter (*D*):

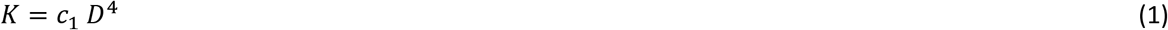

The symbol *C_1_* in eq. 1, and *C_2_…c_6_* in the further equations, denote constants that do not matter for our final results.

Xylem safety can be described by a vulnerability function (*P*), which describes how sapwood conductivity declines with negative xylem water potential *ψ*. Among commonly used xylem vulnerability functions, the Weibull function (eq. 2) has the advantages that it always approaches 1 as *ψ* → 0 and that its parameter *a* does not vary significantly with *ψ_50_* (Duursma & Choat, 2017).

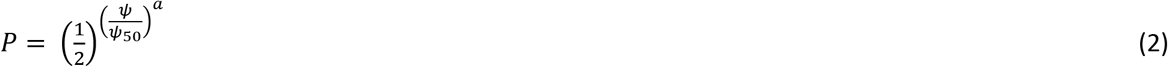

In eq.2, *ψ_50_* is *ψ* that causes 50% loss of conductivity, which is a genetically determined functional trait (Lamy *et al*., 2014; Lobo *et al*., 2018; Pritzkow *et al*., 2020), evolutionarily adapted to the environmental conditions experienced by a species. The parameter *a* controls the shape of *P*, and was estimated based on relationship between *ψ_50_* and *ψ_88_* for each species using the data in (Choat *et al*., 2012) (Appendix). For angiosperms mean *a*=2.48 ± 0.15 (n=153) and for gymnosperms mean *a*=5.34 ± 0.40 (n=29), where ± denotes SE.

The relative importance of different underlying traits in determining *ψ_50_* varies among studies and species, e.g. pit features and vessel length being most important in Acer species (Lens *et al*., 2011) while *D* and vessel grouping are important in other species (Scholz *et al*., 2013; Levionnois *et al*., 2021). Across species, there is an inverse relationship between safety and *D* (eq. 3, where *d* < 0), which is strong in some studies e.g. (Sperry *et al*., 2006; Hacke *et al*., 2015) and weaker, but significant, across multiple studies combined (Hacke *et al*., 2017). Thus, although xylem conductivity and safety are determined by different traits in different species, across species *D* emerges as a reasonable proxy for conduit-level conductivity and safety. From an eco-evolutionary standpoint, a coordination between *D* and other xylem traits should be expected in order to avoid inefficiencies and bottlenecks in the conductive system as a whole. Furthermore, *D* is not only linked to xylem conductivity and safety, but also influences conduit wall thickness and the construction cost of conduits, which all together suggests that *D* should be subject to a strong selection pressure (Koçillari *et al*., 2021).

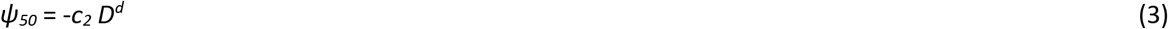

We estimated *d* in eq. 3 based on the data in (Sperry *et al*., 2006) using linear regression of ln(*ψ_50_*) versus ln(*D*). For angiosperms *d* = −1.20 ± 0.14, r^2^ = 0.72 (n=29) and for gymnosperms *d* = −1.32 ±0.63, r^2^ = 0.39 (n=18). Although other datasets may yield slightly different values of *d*, e.g. mean *d* = −0.77 in angiosperms for (Hacke *et al*., 2017), this would only have minor quantitative effects on the results as long as *d*< 0.

Because both *K* and *ψ_50_* depend on *D*, eqs. 1 and 3 can be combined into eq. 4, which represents a conduit-level tradeoff between conductive capacity (*K*) and safety (*ψ_50_*).

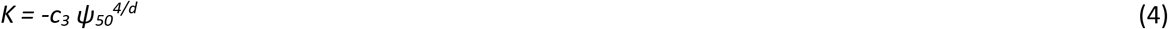

This tradeoff (eq. 4) is defined at the level of conductive tissue (or equivalently, probabilistically at the individual-conduit level) rather than for whole-sapwood tissue. We emphasize that whole-sapwood conductance also depends on the number of conduits per unit sapwood area which weakens this tradeoff at the whole-plant level (Fig. 1).

**Fig. 1.**
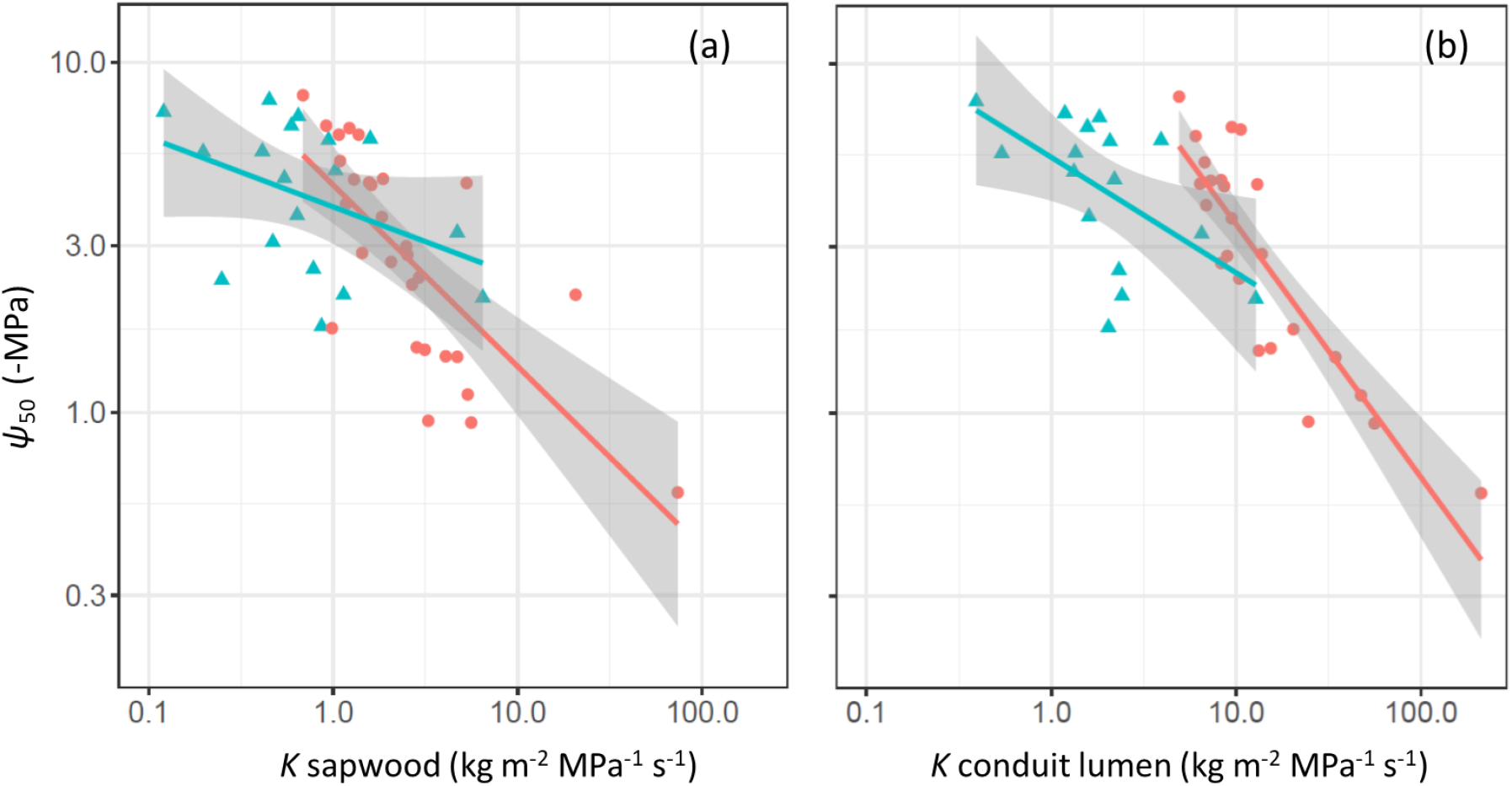
Tradeoff between xylem tolerance to negative water potential (*ψ_50_*) and maximal conductive capacity (*K*). Panels show measurements for whole sapwood (a) and for conduits only (b). Points show observations of angiosperms (red circles) and gymnosperms (blue triangles) and lines show SMA regressions with SE bands, where r^2^= 0.54, 0.16 for sapwood (a) and r^2^= 0.76, 0.32 for conduits (b), for angiosperms (n=29) and gymnosperms (n=18), respectively. Data from (Sperry *et al*., 2006).

Measured vulnerability curves represent the instantaneous loss of conductivity (cavitation) but do not say anything about its reversibility and long-term costs. Conductivity losses are not always reversible (Anderegg *et al*., 2013; Pellizzari *et al*., 2016), which means that consecutive loss events multiply. In this case the total remaining conductivity (*P*_t_) can be described as:

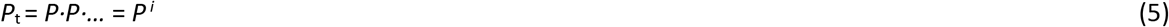

In eq 5, *i* determines the degree of loss accumulation (irreversibility). If *i*= 1, there is no accumulation of effects; If *i*= 2, the fitness-costs of the accumulated loss of conductivity is equal in magnitude to the short-term effects; If *i* > 2, the accumulated (long-term) effects are larger than the instantaneous effects.

### Optimal adaptation of conduit efficiency and safety

The higher the conductance of water through the xylem, the larger the stomatal conductance and carbon uptake can be. Thus, everything else being equal, higher xylem conductance increases fitness. Further, to make the most of acquired resources, all plant organs should be constructed with maximal efficiency, i.e., their function (or fitness contribution) per resources invested should be maximized. The benefit of the xylem is water transport and the costs accrue from the structural investment in fibers and lignified tissues required to build xylem walls (Sperry, 2003). Thus, optimal xylem conduits should have a high conductivity (*K*) per unit mass (*M*), i.e. conductive efficiency (*K*/*M*).

Conduit mass depends on diameter, wall thickness (*T*), and wall tissue density (*ρ*). Biophysical considerations, supported by empirical observations (Hacke *et al*., 2001; Sperry *et al*., 2006), indicate that for conduit walls to withstand negative pressure without imploding, the ratio of conduit wall thickness to conduit diameter (*T/D*) scale with *ψ_50_* according to eq.6 (Hacke *et al*., 2001).

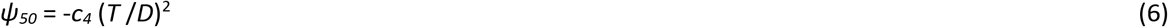

The mass per unit length of a conduit (*M*, eq. 7) can be approximated (by neglecting the difference between outer and inner conduit diameter) by *π·D·T·ρ* and be expressed in terms of *ψ_50_* using eqs. 6 and 3. The approximation introduces only a minor error in optimal *ψ_50_* at highly negative values of *ψ_min_* (1.5% at *ψ_min_* −10 MPa).

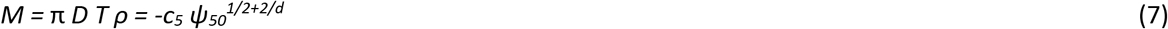

In addition to having a high conductivity per mass, optimal conduits should tolerate low *ψ* with minimal risk of conductivity loss (cavitation), i.e. *P* (eq. 2) should be as large as possible at the minimum operating water potential (*ψ_min_*, the lowest *ψ* measured for a given site and species). Since *ψ_min_* reflects the integrated effect of water flux constraints due to environmental factors and plant stomatal behaviour, it exerts a strong selective force on the xylem (Bhaskar & Ackerly, 2006). Combining the two criteria for optimal xylem function - high conductive efficiency and high tolerance to low *ψ* – in the most parsimonious way, we obtain the optimality criterion, or fitness proxy, 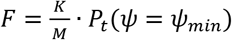. Based on eqs. 2, 4, 5 and 7, *F* can be expressed as a function of *ψ_50_* (eq.8).

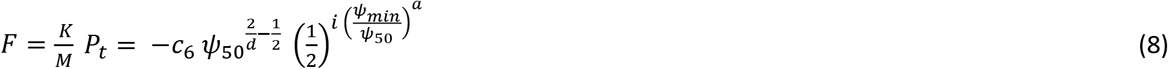

The optimal *ψ_50_* results from the tradeoff between increased conductive capacity per unit conduit biomass and increased vulnerability with increasing *ψ_50_* (Supplementary Fig. S1). *F* is maximized when d*F*/d*ψ_50_* = 0, which results in an optimal *ψ*_50_* that is proportional to *ψ_min_* (eq. 9, derivation in Appendix).

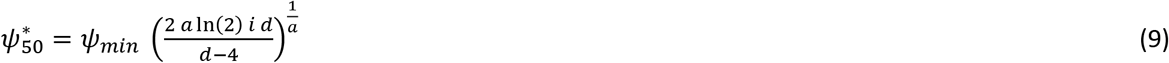

In agreement with observations, eq. 9 implies a linear relationship between *ψ_min_* and *ψ_50_* with zero intercept, i.e. a constant ratio, or safety factor *ψ_50_/ψ_min_* (Fig. 2). For angiosperms, the observed mean safety factor (0.92) is almost identical to the predicted (0.91) for the model without accumulation of conductivity losses (*i*= 1). For gymnosperms, the observed safety factor is 1.7, which can be explained by the model with conductivity loss accumulation (*i* ≠ 1). For this model, *i* was estimated based on the observations, resulting in *i* = 0.99 ± 0.057 (not significant different from 1) for angiosperms and *i* = 9.28 ± 1.94 for gymnosperms, indicating that gymnosperms are much more sensitive to conductivity loss accumulation than angiosperms. The difference in *i* was the main explanation for the higher safety factor in gymnosperms than in angiosperms, whereas the slightly steeper conductivity-safety tradeoff (lower *d*) and the steeper slope of vulnerability function (due to larger *a*) explained 22% of the effect. In addition to the interspecies variation, the model also explains the increase in negative *ψ_50_* with *ψ_min_* along the hydraulic path from roots to branches in two gymnosperm species (Fig. 2c).

**Fig. 2.**
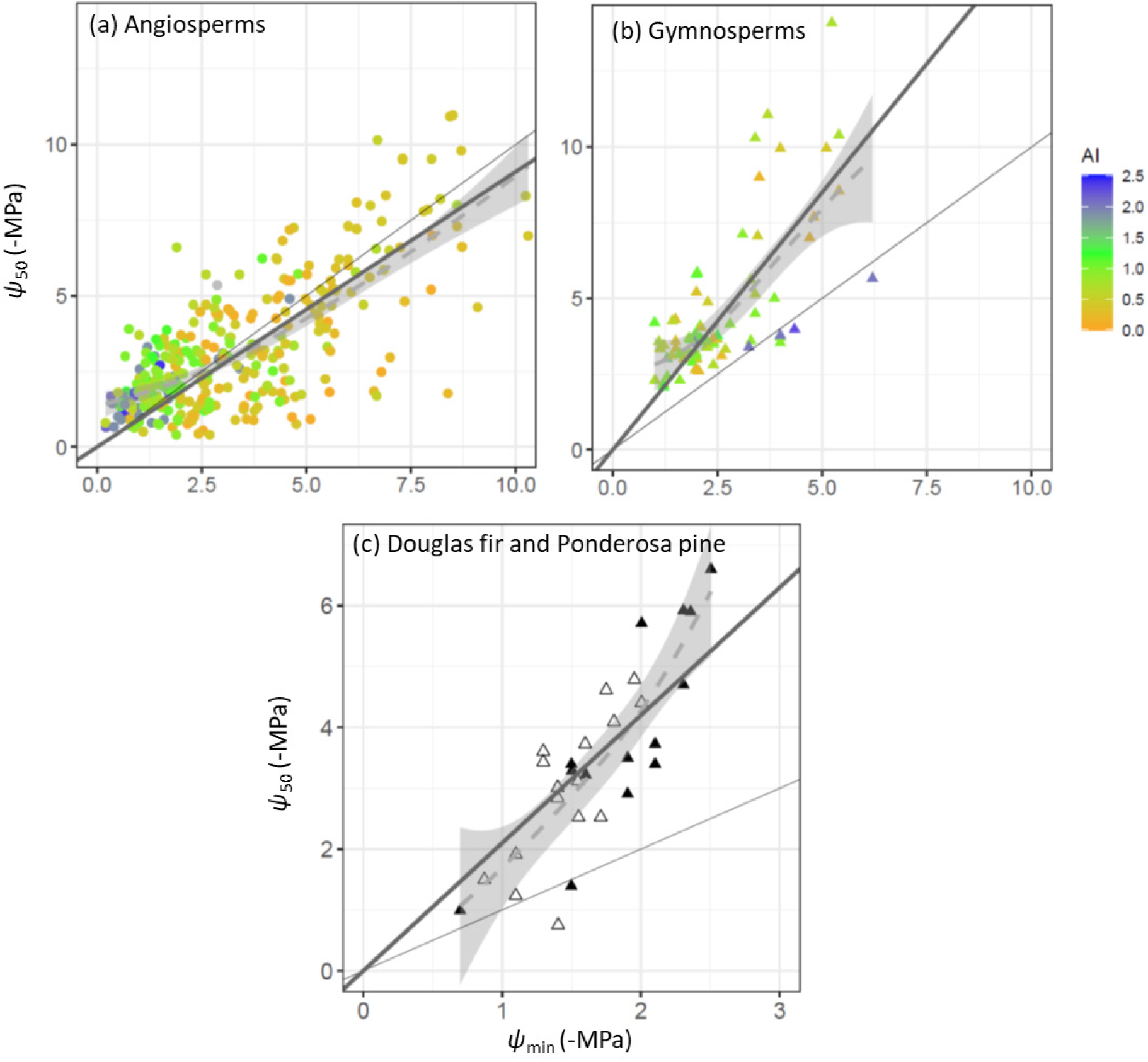
Observed and modelled xylem tolerance to negative water potential (*ψ*_50_) versus minimum plant water potential (*ψ*_min_). (a and b) Measurements in terminal branches for different species and sites (Liu *et al*., 2019), where colors indicate site aridity index (AI) from arid (orange) to wet (blue). Symbol shape indicates angiosperms (a, circles) and gymnosperms (b, triangles). (c) Measurements (means of 5-6 individuals) along the hydraulic path from roots, trunks and branches in Douglas fir (closed triangles) and Ponderosa pine (open triangles) (Domec *et al*., 2009). The dashed lines with shading show smoothed mean and SE intervals of the observed relationships. The straight thick lines are the model predictions (eq. 9). The thin black line is the 1:1 line. Modeled versus observed r^2^ was 0.64 for Douglas fir and Ponderosa pine (n=30), 0.51 for angiosperms (n=338), and 0.48 for gymnosperms (n = 83).

Importantly, the proposed optimality principle defines the intrinsic properties of conduit tissue but does not determine whole sapwood or whole stem conductance. In a whole plant perspective, the principle of optimal conductive efficiency operates at the lowest organizational level, i.e. the level of conduit tissue. Higher-level organizational principles of xylem function are not alternative but complementary to our tissue level principle. While other principles control the variation in xylem structure with plant height or variation in behavioral strategies, optimal conduit efficiency and safety is always maintained at the conduit level. For example, while our principle determines ***ψ*_50_** and *D* at the top of stem, the widened pipe model predicts the relative increase of *D* and decrease of ***ψ*_50_** towards the base (Koçillari *et al*., 2021).

## Discussion

It was previously not explained why so many plants operate with such a small safety margin (*ψ_50_ - ψ_min_*). Our model explains why this is the case – it is the optimal combination of conductive capacity per biomass (*K/M*) and tolerance to the lowest *ψ* experienced by a plant in its environment. The variation in conductive capacity (*K*) relative to low *ψ* tolerance is constrained by an intrinsic physiological tradeoff linked to conduit diameter (*D*). Importantly, this tradeoff is most relevant at the level of conductive tissue only rather than for the whole sapwood (Fig. 1). Weak tradeoffs observed at the whole-sapwood level (Gleason *et al*., 2016) may be caused by the large variability among species in the number of conduits per sapwood area (Zanne *et al*., 2010) and the associated variation in conductivity. Although vulnerability is often more strongly linked to pit- and other traits than to *D* within groups of similar species, across a wide range of species *D* provides a proxy for vulnerability that allows us to link it to xylem conductivity and construction costs. The relevance of optimal adaptation of *D* based on a conductive efficiency – safety tradeoff is further supported by its success in predicting the relative tip to base widening of *D* across species (Koçillari *et al*., 2021).

The model further provides a theoretical explanation and quantification of the previously observed general increase in the hydraulic safety margin with increasingly negative *ψ_min_* (Meinzer *et al*., 2009; Choat *et al*., 2012), which results in a constant safety factor (*ψ_50_ / ψ_min_*) across species of the same type (Fig. 2). Although relevant observations of variation in *ψ_min_* and *ψ_50_* within individuals are yet limited, the available data suggest that the same optimality principle determines *ψ_50_* both across different species and sites and along the xylem flow path within individuals (Fig. 2).

Our simple theoretical model is limited by the few traits explicitly accounted for, not including variation in capacitance, vessel grouping, and pit structure. However, because consistent data on such additional traits are not widely available across species, adding more traits in the model is unlikely to improve its performance, already explaining 50% of the variation in *ψ_50_* across species. Whereas the model captures well the overall trends in *ψ_50_, ψ_50_* for less negative *ψ_min_* (wet sites) is more negative than predicted by the model (Fig. 2a, b). This bias could be related to an under-estimation of negative *ψ_min_* at wetter sites owing to lower sampling of rare dry days at such sites compared to more frequently occurring dry days at drier sites. Plants at wetter sites may be adapted to infrequent drought events which may be missed in the sampling of *ψ_min_* (Martinez Vilalta *et al*., 2021).

The model accurately predicts the observed mean safety factor *ψ_50_ / ψ_min_* = 0.92 in angiosperms, without including irreversible or accumulating loss of conductivity. This may be related to the ability of angiosperms to actively reverse embolism due to the presence of sieve tubes and companion cells in their parenchyma (Johnson *et al*., 2012; Kiorapostolou *et al*., 2019). The much higher mean safety factor in gymnosperm *ψ_50_ / ψ_min_* = 1.7 indicates an adaptation to accumulating effects of conductivity loss and a low reversal capacity (high *i*). A negative relationship between safety factor and embolism reversal capacity is supported by previous observations (Ogasa *et al*., 2013). The importance of irreversible effects of conductivity loss for gymnosperms is further supported by the observations of a low lethal negative *ψ* close to *ψ_50_* (Liang *et al*., 2021).

The invariant *ψ_50_* / *ψ_min_* ratio predicted by our theory suggests that the presence of larger conduit diameters in taller plants (Olson *et al*., 2018) does not necessarily mean that they operate at a higher risk compared to shorter plants. Rather, our hypothesis suggests that taller plants with higher *ψ_50_* should also have higher *ψ_min_* than shorter plants, which is in agreement with observations across a large number of species and environments (Liu *et al*., 2019). In a growing tree with increasing conduit diameter, optimal *ψ_50_* / *ψ_min_* ratio could be maintained by increasing whole xylem conductivity through additional conduits, more conservative stomatal regulation, or by means of increased water uptake with deeper roots, i.e. drought avoidance (Brum *et al*., 2017; Oliveira *et al*., 2021). Nevertheless, if these compensatory mechanisms are hampered by severe drought, taller trees would still suffer more than shorter trees (Rowland *et al*., 2015).

In conclusion, our results show that apparent risky hydraulic behavior of plants can be explained by an eco-evolutionarily optimal design of xylem conduits, constrained by a strong xylem efficiency – safety tradeoff at the scale of individual conduits. *ψ_50_* is universally proportional to *ψ_min_*, corresponding to a constant mean safety factor *ψ_50_* / *ψ_min_* ≈ 0.9 and 1.7 for angiosperms and gymnosperms, respectively. The large safety factor in gymnosperms is likely an adaptation to their small capacity to recover from loss of conductivity compared to angiosperms. The constant safety factor holds across environments and species, and potentially also within individual trees, and thus provides a powerful principle for simplifying and improving plant and vegetation models.

## Acknowledgements

OF and PF were supported by Knut and Alice Wallenberg foundation (Grant 2018.0259). JJ was supported by the European Commission through a Marie Skłodowska-Curie Actions fellowship (Grant No. 841283 – Plant-FATE). OF, JJ, and FH gratefully acknowledge funding from the International Institute for Applied Systems Analysis (IIASA) and the National Member Organizations that support the institute.

## Author contributions

O.F. conceived the original idea, analyzed the data, and wrote the draft manuscript. J.J. and P.F. contributed complementary mathematical analyses. F.H. and J.J. provided complementary data. All authors contributed to conceptual development and the final manuscript.

## Appendix Mathematical derivations

### Derivation of optimal *ψ_50_*

The combined conductive efficiency and safety (*F*) is given by eq.8, as illustrated in Fig. A1.

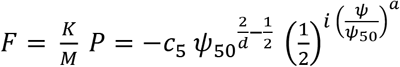

Maximization of *F* with respect to ψ_50_ implies that 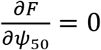 and that 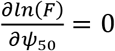.

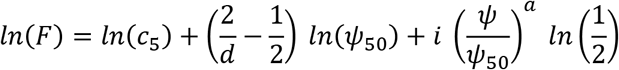

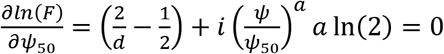, which is solved for *Ψ*_50_ to yield optimal *Ψ*_50_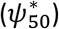 as a function of *Ψ* = *Ψ_min_*:

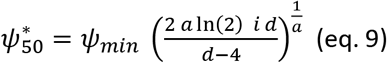

**Fig A1.**
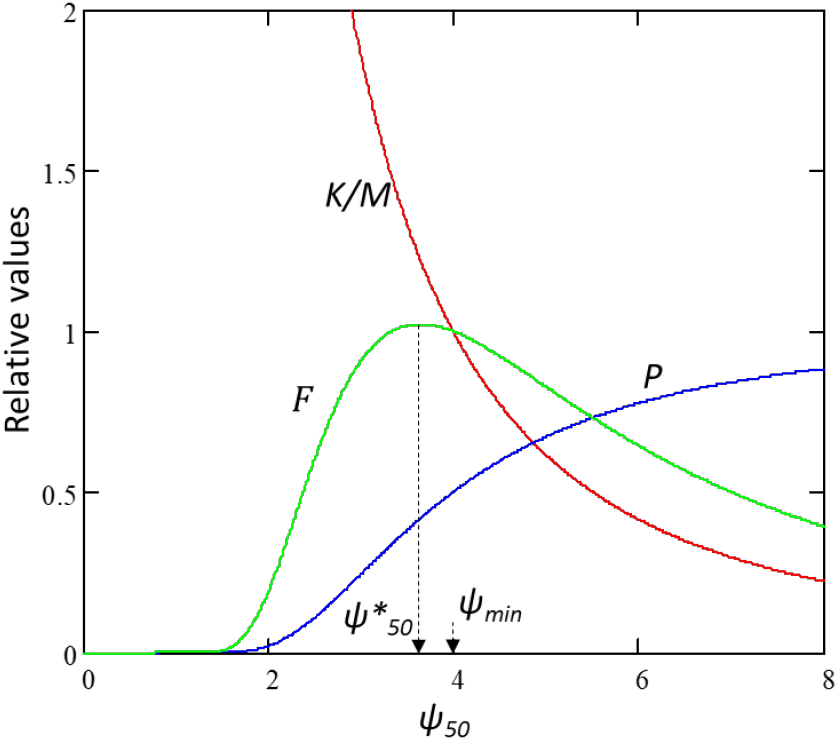
Optimal *ψ*_50_* resulting from maximization of 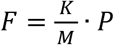 (eq. 8), where 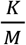(red line) and *F* (green line) are shown normalized with respect to their values at *ψ_50_*=*ψ_min_*.

### Calculation of the parameter *a* in the vulnerability function

*Ψ_88_* corresponds to 88% loss of conductivity, which is inserted into the vulnerability function *P* (eq. 2), which is solved for a.

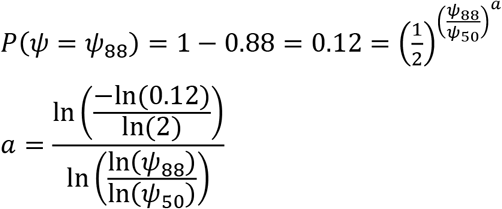

## References

Anderegg WRL, Plavcová L, Anderegg LDL, Hacke UG, Berry JA, Field CB. 2013. Drought’s legacy: multiyear hydraulic deterioration underlies widespread aspen forest die-off and portends increased future risk. Global Change Biology 19(4): 1188–1196.

Anderegg WRL, Wolf A, Arango-Velez A, Choat B, Chmura DJ, Jansen S, Kolb T, Li S, Meinzer FC, Pita P, Resco de Dios V, Sperry JS, Wolfe BT, Pacala S. 2018. Woody plants optimise stomatal behaviour relative to hydraulic risk. Ecology Letters 21(7): 968–977.

Bhaskar R, Ackerly DD. 2006. Ecological relevance of minimum seasonal water potentials. Physiologia Plantarum 127(3): 353–359.

Brum M, Teodoro G, Abrahão A, Oliveira R. 2017. Coordination of rooting depth and leaf hydraulic traits defines drought-related strategies in the campos rupestres, a tropical montane biodiversity hotspot. Plant and Soil 420: 1–14.

Choat B, Jansen S, Brodribb TJ, Cochard H, Delzon S, Bhaskar R, Bucci SJ, Feild TS, Gleason SM, Hacke UG, Jacobsen AL, Lens F, Maherali H, Martínez-Vilalta J, Mayr S, Mencuccini M, Mitchell PJ, Nardini A, Pittermann J, Pratt RB, Sperry JS, Westoby M, Wright IJ, Zanne AE. 2012. Global convergence in the vulnerability of forests to drought. Nature 491(7426): 752–755.

Deans RM, Brodribb TJ, Busch FA, Farquhar GD. 2020. Optimization can provide the fundamental link between leaf photosynthesis, gas exchange and water relations. Nature Plants 6(9): 1116–1125.

Domec J-C, Warren JM, Meinzer FC, Lachenbruch B. 2009. Safety Factors for Xylem Failure by Implosion and Air-Seeding Within Roots, Trunks and Branches of Young and Old Conifer Trees. IAWA Journal 30(2): 101–120.

Duursma RA, Choat B. 2017. fitplc: an R package to fit hydraulic vulnerability curves. Journal of Plant Hydraulics 4: e002.

Franklin O, Harrison SP, Dewar R, Farrior CE, Brännström Å, Dieckmann U, Pietsch S, Falster D, Cramer W, Loreau M, Wang H, Mäkelä A, Rebel KT, Meron E, Schymanski SJ, Rovenskaya E, Stocker BD, Zaehle S, Manzoni S, van Oijen M, Wright IJ, Ciais P, van Bodegom PM, Peñuelas J, Hofhansl F, Terrer C, Soudzilovskaia NA, Midgley G, Prentice IC. 2020. Organizing principles for vegetation dynamics. Nature Plants 6(5): 444–453.

Gleason SM, Westoby M, Jansen S, Choat B, Hacke UG, Pratt RB, Bhaskar R, Brodribb TJ, Bucci SJ, Cao K-F, Cochard H, Delzon S, Domec J-C, Fan Z-X, Feild TS, Jacobsen AL, Johnson DM, Lens F, Maherali H, Martínez-Vilalta J, Mayr S, McCulloh KA, Mencuccini M, Mitchell PJ, Morris H, Nardini A, Pittermann J, Plavcová L, Schreiber SG, Sperry JS, Wright IJ, Zanne AE. 2016. Weak tradeoff between xylem safety and xylem-specific hydraulic efficiency across the world’s woody plant species. New Phytologist 209(1): 123–136.

Hacke UG, Sperry JS, Pockman WT, Davis SD, McCulloh KA. 2001. Trends in wood density and structure are linked to prevention of xylem implosion by negative pressure. Oecologia 126(4): 457–461.

Hacke UG, Spicer R, Schreiber SG, Plavcová L. 2017. An ecophysiological and developmental perspective on variation in vessel diameter. Plant, Cell & Environment 40(6): 831–845.

Hacke UG, Venturas MD, MacKinnon ED, Jacobsen AL, Sperry JS, Pratt RB. 2015. The standard centrifuge method accurately measures vulnerability curves of long-vesselled olive stems. New Phytologist 205(1): 116–127.

Hölttä T, Mencuccini M, Nikinmaa E. 2011. A carbon cost–gain model explains the observed patterns of xylem safety and efficiency. Plant, Cell & Environment 34(11): 1819–1834.

Johnson DM, McCulloh KA, Woodruff DR, Meinzer FC. 2012. Hydraulic safety margins and embolism reversal in stems and leaves: Why are conifers and angiosperms so different? Plant Science 195: 48–53.

Kiorapostolou N, Da Sois L, Petruzzellis F, Savi T, Trifilò P, Nardini A, Petit G. 2019. Vulnerability to xylem embolism correlates to wood parenchyma fraction in angiosperms but not in gymnosperms. Tree Physiology 39(10): 1675–1684.

Koçillari L, Olson ME, Suweis S, Rocha RP, Lovison A, Cardin F, Dawson TE, Echeverría A, Fajardo A, Lechthaler S, Martínez-Pérez C, Marcati CR, Chung K-F, Rosell JA, Segovia-Rivas A, Williams CB, Petrone-Mendoza E, Rinaldo A, Anfodillo T, Banavar JR, Maritan A. 2021. The Widened Pipe Model of plant hydraulic evolution. Proceedings of the National Academy of Sciences 118(22): e2100314118.

Lamy J-B, Delzon S, Bouche PS, Alia R, Vendramin GG, Cochard H, Plomion C. 2014. Limited genetic variability and phenotypic plasticity detected for cavitation resistance in a Mediterranean pine. New Phytologist 201(3): 874–886.

Lens F, Sperry JS, Christman MA, Choat B, Rabaey D, Jansen S. 2011. Testing hypotheses that link wood anatomy to cavitation resistance and hydraulic conductivity in the genus Acer. New Phytologist 190(3): 709–723.

Levionnois S, Jansen S, Wandji RT, Beauchêne J, Ziegler C, Coste S, Stahl C, Delzon S, Authier L, Heuret P. 2021. Linking drought-induced xylem embolism resistance to wood anatomical traits in Neotropical trees. New Phytologist 229(3): 1453–1466.

Liang X, Ye Q, Liu H, Brodribb TJ. 2021. Wood density predicts mortality threshold for diverse trees. New Phytologist 229(6): 3053–3057.

Liu H, Gleason SM, Hao G, Hua L, He P, Goldstein G, Ye Q. 2019. Hydraulic traits are coordinated with maximum plant height at the global scale. Science Advances 5(2): eaav1332.

Liu H, Ye Q, Gleason SM, He P, Yin D. 2021. Weak tradeoff between xylem hydraulic efficiency and safety: climatic seasonality matters. New Phytologist 229(3): 1440–1452.

Lobo A, Torres-Ruiz JM, Burlett R, Lemaire C, Parise C, Francioni C, Truffaut L, Tomášková I, Hansen JK, Kjær ED, Kremer A, Delzon S. 2018. Assessing inter- and intraspecific variability of xylem vulnerability to embolism in oaks. Forest Ecology and Management 424: 53–61.

Manzoni S, Vico G, Katul G, Palmroth S, Jackson RB, Porporato A. 2013. Hydraulic limits on maximum plant transpiration and the emergence of the safety–efficiency trade-off. New Phytologist 198(1): 169–178.

Martinez Vilalta J, Santiago L, Poyatos R, Badiella L, De Cáceres M, Aranda I, Delzon S, Vilagrosa A, Mencuccini M. 2021. Towards a statistically robust determination of minimum water potential and hydraulic risk in plants. New Phytologist 232(1): 404–417.

McCulloh KA, Sperry JS, Adler FR. 2003. Water transport in plants obeys Murray’s law. Nature 421(6926): 939–942.

Meinzer FC, Johnson DM, Lachenbruch B, McCulloh KA, Woodruff DR. 2009. Xylem hydraulic safety margins in woody plants: coordination of stomatal control of xylem tension with hydraulic capacitance. Functional Ecology 23(5): 922–930.

Ogasa M, Miki NH, Murakami Y, Yoshikawa K. 2013. Recovery performance in xylem hydraulic conductivity is correlated with cavitation resistance for temperate deciduous tree species. Tree Physiology 33(4): 335–344.

Oliveira RS, Eller CB, Barros FdV, Hirota M, Brum M, Bittencourt P. 2021. Linking plant hydraulics and the fast–slow continuum to understand resilience to drought in tropical ecosystems. New Phytologist 230(3): 904–923.

Olson ME, Soriano D, Rosell JA, Anfodillo T, Donoghue MJ, Edwards EJ, León-Gómez C, Dawson T, Camarero Martínez JJ, Castorena M, Echeverría A, Espinosa CI, Fajardo A, Gazol A, Isnard S, Lima RS, Marcati CR, Méndez-Alonzo R. 2018. Plant height and hydraulic vulnerability to drought and cold. Proceedings of the National Academy of Sciences 115(29): 7551–7556.

Pellizzari E, Camarero JJ, Gazol A, Sangüesa-Barreda G, Carrer M. 2016. Wood anatomy and carbon-isotope discrimination support long-term hydraulic deterioration as a major cause of drought-induced dieback. Global Change Biology 22(6): 2125–2137.

Pritzkow C, Williamson V, Szota C, Trouvé R, Arndt SK. 2020. Phenotypic plasticity and genetic adaptation of functional traits influences intra-specific variation in hydraulic efficiency and safety. Tree Physiology 40(2): 215–229.

Rowland L, da Costa ACL, Galbraith DR, Oliveira RS, Binks OJ, Oliveira AAR, Pullen AM, Doughty CE, Metcalfe DB, Vasconcelos SS, Ferreira LV, Malhi Y, Grace J, Mencuccini M, Meir P. 2015. Death from drought in tropical forests is triggered by hydraulics not carbon starvation. Nature 528(7580): 119–122.

Sanchez-Martinez P, Martínez-Vilalta J, Dexter KG, Segovia RA, Mencuccini M. 2020. Adaptation and coordinated evolution of plant hydraulic traits. Ecology Letters 23(11): 1599–1610.

Savage VM, Bentley LP, Enquist BJ, Sperry JS, Smith DD, Reich PB, Von Allmen EI. 2010. Hydraulic trade-offs and space filling enable better predictions of vascular structure and function in plants. Proceedings of the National Academy of Sciences of the United States of America 107(52): 22722–22727.

Scholz A, Rabaey D, Stein A, Cochard H, Smets E, Jansen S. 2013. The evolution and function of vessel and pit characters with respect to cavitation resistance across 10 Prunus species. Tree Physiology 33(7): 684–694.

Sperry JS. 2003. Evolution of Water Transport and Xylem Structure. International Journal of Plant Sciences 164(S3): 115–127.

Sperry JS, Hacke UG, Pittermann J. 2006. Size and function in conifer tracheids and angiosperm vessels. American Journal of Botany 93(10): 1490–1500.

Venturas MD, Sperry JS, Hacke UG. 2017. Plant xylem hydraulics: What we understand, current research, and future challenges. Journal of Integrative Plant Biology 59(6): 356–389.

Wang H, Prentice IC, Keenan TF, Davis TW, Wright IJ, Cornwell WK, Evans BJ, Peng C. 2017. Towards a universal model for carbon dioxide uptake by plants. Nature Plants 3(9): 734–741.

Wang Y, Sperry JS, Anderegg WRL, Venturas MD, Trugman AT. 2020. A theoretical and empirical assessment of stomatal optimization modeling. New Phytologist 227(2): 311–325.

West GB, Brown JH, Enquist BJ. 1999. A general model for the structure and allometry of plant vascular systems. Nature 400(6745): 664–667.

Wolf A, Anderegg WRL, Pacala SW. 2016. Optimal stomatal behavior with competition for water and risk of hydraulic impairment. Proceedings of the National Academy of Sciences 113(46): E7222–E7230.

Zanne AE, Westoby M, Falster DS, Ackerly DD, Loarie SR, Arnold SEJ, Coomes DA. 2010. Angiosperm wood structure: Global patterns in vessel anatomy and their relation to wood density and potential conductivity. American Journal of Botany 97(2): 207–215.

